# Fabrication of the high-resistance patch-clamp pipettes for mitochondrial electrophysiological studies using optimized two-step method

**DOI:** 10.64898/2026.05.05.723071

**Authors:** Noor Mohamed, Oleg Artemchuk, Sasan Rabieh, Pablo Peixoto, Tim Bromage, Evgeny Pavlov

## Abstract

The patch-clamp experimental technique is widely used to study the electrical properties of ion channels in biological and artificial lipid membranes. The key to the high quality of the experiments is the manufacturing of glass pipettes that provide highly electrically resistant contact between the edge of the pipette tip and the lipid bilayer. Preparation of the pipettes is particularly challenging for studies of the mitochondrial membranes due to the need for very small pipette tip sizes. Here, we present a robust procedure for producing pipettes suitable for experiments with native mitochondrial membranes. This procedure involves a two-step approach: initial fabrication of relatively large glass micropipettes using a standard micropipette puller, followed by tip refinement using a microforger to achieve smooth glass surface and reduced opening size. Pipette tip diameters and surface structure were examined using field emission – scanning electron microscopy (FE-SEM) imaging to assess the effects of variable parameters on pipette geometry and size. The resulting pipettes were validated in patch-clamp recording of the mitochondrial inner membranes. This approach enables the reproducible production of optimized pipettes for mitochondrial patch-clamp experiments, improving the quality and throughput of electrophysiological recordings of the mitochondrial ion channels.

## Introduction

Mitochondrial ion channels play critical roles in the modulation of cellular energy production, calcium signaling, and cell death, and patch clamp studies have provided unique insights into ion transport and mitochondrial membrane protein function^1-5^. Patch-clamp technique is an essential tool in mitochondrial physiology and pathology studies, allowing researchers to investigate the electrical properties of mitochondrial ion channels. However, the application of patch-clamp approaches to mitochondrial membranes is technically demanding, requiring micropipettes with extremely fine tips to form a high-resistance seal with small and delicate mitochondrial membranes.

A major challenge in the field is the absence of standardized protocols for fabricating micropipettes for mitochondrial patch clamping with consistent size and geometry. Pipettes for mitochondrial studies require exceptionally small tip sizes compared to those used in conventional whole-cell or cell-excised patch configurations. Currently, fabrication of such pipettes relies largely on empirical optimization of the standard pipette puller parameters through trial-and-error approaches. Some practical considerations and challenges have been described in details in recent work from the Bertholet lab^6^. However, specific optimized parameters generating suitable small pipettes strongly depend on environmental factors (e.g. air humidity) and require continuous adjustment. As a result, generating functional micropipettes is frequently inconsistent, difficult to reproduce, and difficult to replicate across studies. This limitation is particularly critical in mitochondrial membranes studies, where poor seal quality of the membrane and resulting leak currents may be misinterpreted as endogenous electrical activity.

Here, we aimed to develop a reproducible and optimized protocol for creating micropipettes that consistently meet the unique requirements of mitochondrial patch-clamp experiments. The proposed method employs a two-step process of pipette pulling followed by controlled tip heating, which significantly improves the reliability and reproducibility of mitochondrial electrophysiological recordings.

## Materials and Methods

### Pipette Pulling

Pipettes were fabricated using a Flaming/Brown micropipette puller (Model P-1000, Sutter Instrument). Fire-polished borosilicate glass capillaries with an outer diameter of 1.5 mm and an inner diameter of 0.75 mm (BF150-86-10, Sutter Instrument) were employed. The puller parameters evaluated included heat, pull, and velocity. To assess the impact of each parameter on the diameters of pipette tips, all other parameters were held constant while the parameter of interest was varied. Table 1 outlines the parameter values used in these experiments. The “Heat” parameter value depends on the RAMP value of pipette puller. For all experiments reported here, the RAMP value was 554. Furthermore, ten pipettes were fabricated under identical parameter settings, and their tip diameters were measured using field emission – scanning electron microscopy (FE-SEM) to assess the reproducibility of the puller. The results are illustrated in Figure 1 A, B.

**Table 1.** Parameter values for pipette pulling.

| **Table 1 Parameter values for pipette pulling.** |  |  |  |
| --- | --- | --- | --- |
| **Experiment** | **Heat** | **Pull** | **Velocity** |
| Effect of Heat | 554, 559, 564, 569, 574 | 0 | 30 |
| Effect of Pull | 564 | 0, 10, 20, 30 | 30 |
| Effect of Velocity | 564 | 0 | 26, 28, 30, 32 |

**Figure 1.**
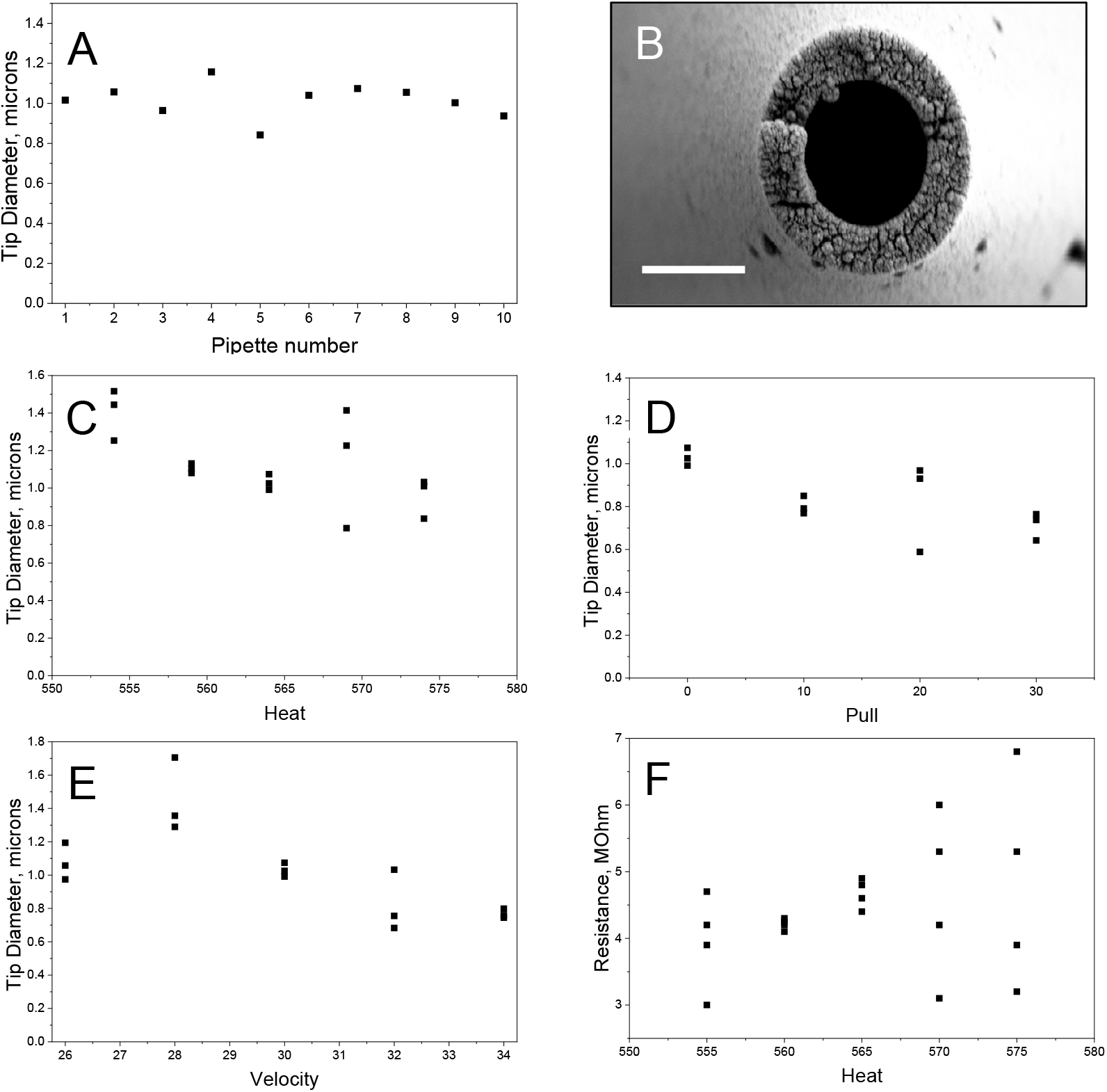
*Optimization of the pipette puller parameters*. **(A)** Pipette pulling records. 10 pipettes were pulled under identical pulling parameters and their corresponding tip sizes were measured using FE-SEM. **(B)** FE-SEM imaging of pipette after pulling (scale bar, 1 micron). **(C)** The effect of heat on tip diameter. Heat is measured in milliamps. Notable changes in heat necessitate an adjustment of at least 5 milliamps. Sample sizes are n = 3 for measurements at 553, 559, and 564 milliamps, and n = 5 for measurements at 596 and 574 milliamps. **(D)** The effect of pull on tip diameter. Notable changes in pull necessitate an adjustment of at least 10 units. Sample sizes are n = 3.**(E)** The effect of velocity on tip diameter. Notable changes in velocity necessitate an adjustment of at least 10 units. Sample sizes are all n = 3 for all samples except that of 28, which has a sample size of n=4. **(F)** Optimal heat for consistent pipette resistances. Pipette resistance was measured across different heat parameters to determine the parameter that yields the most consistent results. Each condition was tested with a sample size of n = 5.

### Pipette forging

Following the pulling step, glass micropipettes were positioned in the Narishige MF-900 microforge using 35x objective. The filament heating was set at 35 units (approx. 550°C), while the metal filament was positioned at a distance of 12 units from the pipette tip (approximately 34 µm; Fig. 2A). The temperature of the forge and the distance between the filament and the pipette tip were kept constant throughout the experiments. The detailed sample procedure for preparing pipettes is described in Results section.

**Figure 2.**
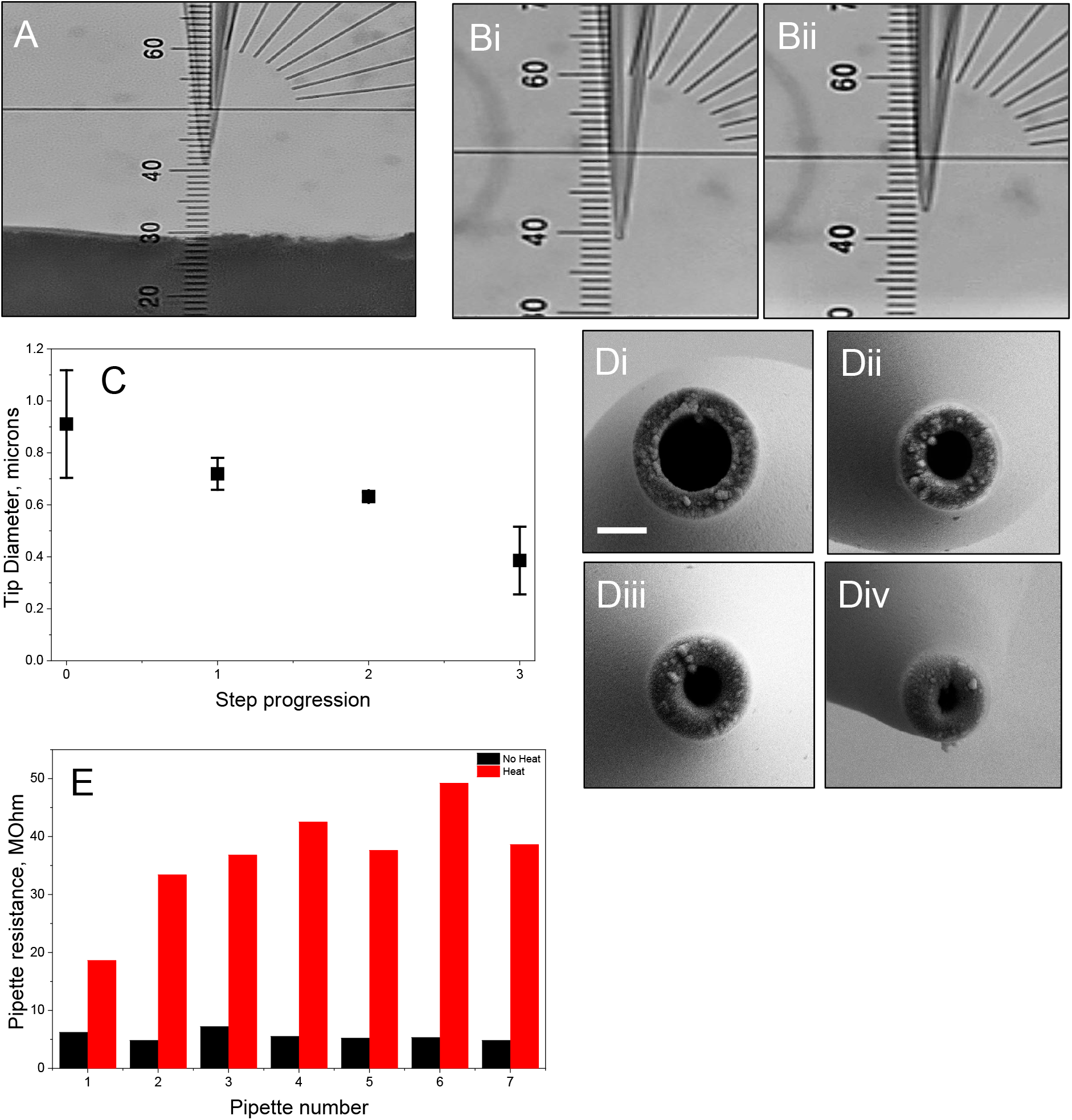
*Optimization of Pipette Tip via Incremental Heat Polishing*. **(A)** Intial setup of the pipette for forging. **(B)** Working distance between pipette and metal filament during forging. Hote the shorted tip after heat application. **(C)** Optimization of pipette tip diameter via incremental heating. Step progression denotes heating steps after the pulling protocol (Step 0 = no heating, Step 1–3 = increasing heating clicks; Step 3 is the optimized protocol). Sample sizes are n = 3. Data are presented as the mean ± SEM. **(D) (Di-Div)** FE-SEM imaging of the optimization of pipette tip diameter via incremental heating. Step progression denotes heating steps after the pulling protocol. **(Di)** pipette pulled at Step 0 parameters. **(Dii)** pipette pulled at Step 1 parameters. **(Diii)** pipette pulled at Step 2 parameters. **(Div)** pipette pulled at Step 3 parameters (optimized protocol). **(E)** Reproducibility of the pipette resistance during repetitive pipette preparation. Note consistent increase in pipette resisance following forging step.

### Field Emission – Scanning Electron Microscopy (FE-SEM) Imaging

Pipette samples were coated with 10 nm of gold using a Safematic compact coating unit (Safematic GmbH, CCU-010, Zizers, Switzerland) to render the surfaces electrically conductive for imaging.

All FE-SEM images were collected using a Zeiss Gemini-300 FE-SEM (Carl Zeiss Microscopy, White Plains, NY, USA) operated in secondary electron emission mode. Images were collected at a field width of 5 µm at high vacuum and an accelerating voltage of 10 kV, a 30 μm aperture, and a working distance of approximately 6.0 mm to characterize pipette tip diameter.

### Mitochondria isolation

All chemicals for mitochondrial isolation and patch-clamp recordings were purchased from Sigma-Aldrich, USA. Fresh mitochondria were isolated from mouse hearts on the day of each experiment using mechanical homogenization followed by differential centrifugation. Hearts were quickly removed from euthanized mice following approval by the Institutional Animal Care and Use Committee. Immediately after dissection, the hearts were rinsed three times in ice-cold isolation buffer composed of 225 mM mannitol, 75 mM sucrose, 5 mM 4-(2-hydroxyethyl)-1-piperazineethanesulfonic acid (HEPES; pH 7.2), and supplemented with 1 mg/mL fatty acid-free bovine serum albumin (BSA).

The rinsed tissue was finely minced using scissors in fresh isolation buffer on ice, then transferred to a 5-mL Teflon-glass homogenizer. Homogenization was performed with 20 gentle strokes of the pestle at 300 rpm while keeping the homogenizer on ice.The resulting homogenate was centrifuged at 800 × g for 10 min at 4°C to remove unbroken tissue and nuclei. The supernatant was carefully collected and subjected to a second centrifugation at 10,000 × g for 10 min at 4°C. The supernatant from this spin was discarded, and the mitochondrial pellet was gently resuspended in 100 μL of sucrose-based buffer containing 225 mM mannitol, 75 mM sucrose, 10 mM EGTA, 5 mM HEPES (pH 7.2).The resuspended mitochondria were kept on ice until further use.

### Mitoplast preparation and patch-clamp recordings

The mitochondrial pellet was resuspended in recording solution (150 mM KCl, 5 mM HEPES, pH 7.2–7.4) supplementaed with mitochondrial respiratory substrates (2 μM rotenone, 5 mM succinate, and 1 mM Na_2_HPO_4_). To induce mitoplast formation, 1 mM CaCl_2_ was added to trigger opening of the mitochondrial permeability transition pore (mPTP) and subsequent swelling. Immediately after calcium addition, the mitochondrial suspension was transferred to a glass-bottom recording chamber.

Calcium-induced swelling and mitoplast formation were monitored in real time under an inverted bright-field microscope (Nikon Eclipse TE300) equipped with a digital camera (Accu-Scope)^7^.Electrical currents were recorded using an Axon 200 amplifier (Molecular Devices) and Clampex software.

## Results and discussion

In our approach, pipette fabrication was done in two steps. Initial step was the preparation of the pipette with the parameters similar to the pipettes used for whole cells patch-clamp. This is a robust procedure with the strategy described in the Sutter manual “The Pipette Cookbook”: https://www.sutter.com/PDFs/pipette_cookbook.pdf^8^. We found that this strategy is highly efficient for production of patch-clamp pipettes up to 10 MOhm in resistance, if measured in standard recoding solution containing 150 mM KCl, 5 mM HEPES, pH 7.2–7.4.

To optimize first (pulling) stage parameters we verified how they affect the pipette size as determined by tip diameter measured using FE-SEM and corresponding pipette resistance. Generally, if parameters remain unchanged pipette size remains stable (Figure 1 A) producing round shaped pipette tip of the diameter around 1 micron (Figure 1B), which corresponded to the pipette resistance of ∼5 MOhm. The pulling parameters, heat, pull, and velocity, had quantifiable effects on pipette tip diameter, (Figure 1, C-E) and resistance (Figure 1, F). Increasing heat levels (measured in milliamps) yielded smaller tip diameters due to enhanced softening of the glass during the pull. Similarly, increasing the velocity parameter produced narrower tips. This likely reflects a faster stretching rate of the molten glass, which elongates the pipette more rapidly and reduces the final diameter. Pull also influenced pipette diameter as higher pull values reduced tip diameter by promoting further elongation of the softened glass.

The second (heat) stage further refined pipette geometry with the goal to make the pipette opening smaller. In this case several parameters needed be optimized: the distance between heating element and pipette tip (Fig. 2 A), the heating temperature and duration and a number to heating applications. The distance and heating temperature were adjusted in such a way to allow the duration of heat to be sufficiently long. We found that applying heat for duration of few seconds provides best reproducibility outcome. It should also be noted that heat application resulted in the shortening of the pipette tip (compare Fig. 2Bi and Bii)

As illustrated in Figure 2C and D, increasing the number of forge heat applications (each lasting 10 seconds) led to progressive reductions in tip diameter. This suggests that post-pulling heat polishing can smoothen and sharpen pipette tips beyond what is achieved by the pulling alone.

Electrical resistance data further supported these observations. As illustrated at figure 2E polishing procedure that significantly reduced pipette opening size resulted in an increase in resistance. Notably, pipettes parameters were highly reproducible.

Following pipettes preparation their utility was tested by patch-clamp measurements of isolated mitoplasts (Fig 3.A). We have found that mitoplasts from the intact mitochondria that were prepared by passive swelling demonstrated low conductance (Fig. 3B, red trace), while mitoplasts prepared by calcium-induced swelling of energized mitochondria demonstrated channel activity which is consistent with the activity of the mitochondrial Permeability Transition Pore (Fig. 3B, black trace). The success rate for obtaining seals was at least 75% with typical conductance of the membranes in the absence of ion channels at the range of 5-10 Gohms.

**Figure 3.**
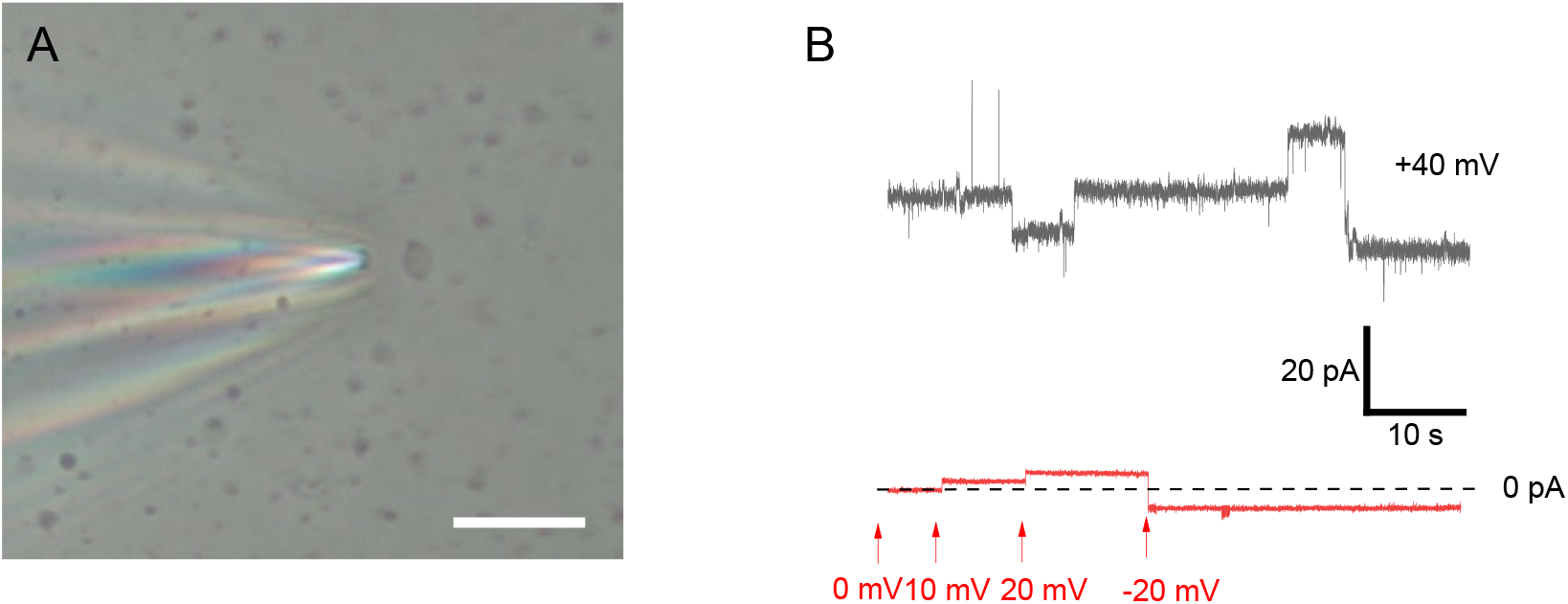
Patch-clamp recodrings of the electrical activity of the mitochondrial inner membrane. **(A).** Image of the mitoplast preparation used in patch-clamp experiments. Mitoplast can be seen as a grey round vesicle. Note that at this magnification size of pathc-clamp pipette can not be estimated. (**B)**. Representative current traces of the the control membrane lacking ion channels activity (red trace) and membrane of calcium treated mitochondria with high-condcutance channel actitity, which is typical for mitochondrial Permeability Transtion Pore.

Taken together, these findings informed the development of a two-stage optimized protocol for the fabrication of patch pipettes tailored for mitochondrial electrophysiology. The protocol combines fine-tuned pulling parameters using a Sutter P-1000 puller, with a heat setting of 564 (RAMP), velocity of 30, and no additional pull force, followed by a three-step heat polishing process using a Narishige MF-900 microforge. This approach reliably produces pipettes with smooth tapers, small inner tip diameters (∼1–2 µm), and resistances between 30–45 MΩ, ideal for forming gigaseals on mitochondrial membranes.

Based on the experimental findings, we developed the following step-by-step instructions to reliably fabricate patch pipettes optimized for mitochondrial electrophysiology. We should note that protocol settings for the puller are for reference, the exact settings will need to be optimized based on the strategies described in the methods and results sections:

### Stage 1: Pulling the Pipette

#### Equipment

- Sutter P-1000 Micropipette Puller
- Borosilicate Glass Capillaries (OD: 1.5 mm, ID: 0.75 mm, length: 10 cm, e.g., BF150-75-10)

#### Procedure

1. Load a clean borosilicate capillary into the Sutter P-1000 puller.
2. Set the puller to the following optimized parameters (these parameters can vary between pullers and specific conditions and should be determined empirically): Heat: 564, Pull: 0, Velocity: 30, Time: 150, Pressure: 500
3. Execute the pull. Discard pipettes that have broken or asymmetrical tapers.
4. Select pipettes with a long, smooth taper and intact, clean tips. Measure resistance (optional at this stage) to verify it falls within the 1–5 MΩ range.

### Stage 2: Heat Polishing the Tip

#### Equipment

1. Narishige MF-900 Microforge with 40x objective
2. Dissecting Microscope or Eyepiece with Bar Scale

#### Procedure

1. Mount the pipette under the microforge microscope.
2. Setup forge heat parameter at 35.
3. Adjust the position so that the pipette tip is approximately 12 grids away from the heated filament (Fig. 2A).
4. Ensure the tip is centered and aligned horizontally with the filament.
5. Perform the following “three-clicks” polishing sequence. Click 1 (Pre-shaping): Gently touch the button to activate heat filament. Hold for 7 seconds, then release. Click 2 (Initial Polishing): After first heat tip becomes shorter (compare figure 2 Bi to Bii) and needs to be repositioned back to the distance of 12 gords, which in case of figure 2Bii will be at 39. Hold heat for 7 seconds, then release. Click 3 (Final Polish): Make final controlled contact. Hold heat for 15 seconds, then release.

### Stage 3: Post-Polishing Inspection and Storage

1. Inspect the tip under a high-magnification microscope.
  - The ideal tip should be smooth, rounded, and free of jagged edges or debris.
  - Target an inner diameter of ∼1–2 µm.
2. Measure resistance again (if applicable); optimal pipettes should show 30–45 MΩ.
3. Store pipettes horizontally in a clean, dust-free environment.
  - Use within 24 hours to avoid contamination and maintain structural integrity.

This study establishes a reproducible and optimized protocol for the fabrication of glass micropipettes suitable for patch-clamp recordings of mitochondrial membranes. By systematically evaluating the effects of pulling parameters, heat, pull, and velocity, on pipette geometry, and refining the heat-polishing process using a microforge, we identified conditions that consistently yield pipettes with tip diameters of approximately 1–2 µm and resistances ranging from 30 to 45 MΩ. Among the tested variables, heat exerted the most significant influence on tip diameter, while incremental heat polishing further improved tip smoothness and resistance consistency. This methodological standardization addresses a major technical limitation in mitochondrial electrophysiology and facilitates the generation of high-quality pipettes necessary for forming stable high-resistance seals. By standardizing micropipette fabrication and reducing reliance on trial-and-error methods, this protocol improves the reproducibility and reliability of mitochondrial patch-clamp recordings. Consistent pipette geometry is essential for achieving stable, high-resistance seals with mitochondrial membranes, which are particularly sensitive to fabrication variability. The ability to generate uniform pipettes enables more accurate and comparable measurements of mitochondrial ion channel activity and membrane properties. This advancement supports broader application of mitochondrial electrophysiology and facilitates more investigation of mitochondrial function in both physiological and disease contexts.

## Notes

### Competing Interest Statement

The authors have declared no competing interest.

### Funding

- National Institute of General Medical Sciences: R35GM139615

